# Molecular insights into the modulation of the 5HT_2A_ receptor by serotonin, psilocin, and the G protein subunit Gqα

**DOI:** 10.1101/2024.07.23.604750

**Authors:** Niklas Viohl, Ali Asghar Hakami Zanjani, Himanshu Khandelia

## Abstract

The 5HT_2A_ receptor (5HT_2A_R) is a G protein-coupled receptor that drives many neuronal functions and is one of the primary targets for psychedelic drugs, which have recently shown promise in treating mental disorders. Understanding ligand interactions and conformational transitions is essential for developing effective pharmaceuticals, but mechanistic details of 5HT_2A_R activation and ligand binding modes remain poorly understood. We conducted all-atom molecular dynamics simulations and free energy calculations of the active and inactive forms of 5HT_2A_R with psilocin and serotonin. Both serotonin and psilocin have higher binding affinities for the orthosteric binding pocket than the extended binding pocket. Active state 5HT_2A_R collapses to a closed state in the absence of Gqα. We also discover a ‘partially-open’ receptor conformation that is intermediate between the active and inactive states. Our discoveries can inform the design of new pharmaceuticals that target specific receptor conformations, potentially leading to more effective treatments for mental disorders.

## INTRODUCTION

Neuronal signal transduction is achieved through an intricate interplay between various neurotransmitters and their corresponding postsynaptic receptors. Neurotransmitter systems in the brain include noradrenergic, dopaminergic, histaminergic, cholinergic, adrenergic, and serotonergic pathways. Among other things, serotonin signaling regulates mood, memory, sexual behavior, sleep, learning, and reward processing and disruption of serotonergic pathways can promote numerous physical and mental disorders like depression, migraine, schizophrenia, and anxiety.^1,2^

One promising group of rediscovered novel antidepressant candidates is psychedelic drugs like psilocybin, MDMA, or LSD. Natural psychedelic compounds like psilocybin or mescaline have been used by various indigenous cultures for centuries, usually integrating them into the framework of ritualistic contexts. Psilocybin occurs in a large variety of fungi known as ‘magic mushrooms’. In the liver, the prodrug psilocybin is rapidly converted into the pharmacologically active psilocin. After the chemical extraction and synthesis of psilocin, LSD, and other psychedelics in the 1950s and 1960s, pharmacological and behavioral research flourished until substance regulations diminished research activity.^3–5^ Several recent clinical studies showed that the administration of hallucinogenic drugs can drastically improve mental disorders, while scarcely causing side effects and adverse reactions.^6–8^

Following the release into the synaptic cleft, serotonin, and other effectors bind to one of the G protein-coupled receptor (GPCR)-type serotonin receptors (5HTRs) and stimulate a transducer-mediated signaling pathway. The binding of serotonin and other stimulants to 5HT_2A_R – the main receptor believed to be responsible for psychedelic and antidepressant effects^5,9^ – induces conformational changes in the intracellular region, that allow for the binding of either β-arrestin or the heterotrimeric Gq protein complex. Ultimately, both transducer pathways alter gene expression patterns and kinase signaling, which in turn stimulate several neuroplastic processes.^10,11^

Due to the comparatively non-specific binding mode, many psychedelics like LSD and psilocin act promiscuously on many neural receptors.^5^ Besides 5HT_2A_R, psilocin, and synthetic psilocin analogs exhibit binding affinities for 5HT_1A_R and 5HT_2C_R^12–14^ and animal studies suggest that psilocin unfolds its psychedelic and antidepressant effects not solely via stimulation of 5HT_2A_R.^15–17^ The structure of 5HT_2C_R in complex with psilocin shows that psilocin is similarly anchored in the orthosteric binding pocket (OBP) via the conserved central D134^3.32^ residue, and the indole core is coordinated through interactions with additional residues in the OBP.^18^ Moreover, psychedelics like ketamine, LSD, and psilocin influence synaptic plasticity, thereby facilitating antidepressant effects by binding to the neurotrophic tyrosine kinase receptor 2 (TrkB) and stimulating endogenous brain-derived neurotrophic factor (BDNF) signaling.^19,20^ It was recently demonstrated that a 5HT_1A_R-selective analog of the psychedelic substance 5-MeO-DMT analog lacks hallucinogenic effects but maintains anxiolytic and antidepressant activities in socially defeated mice.^21^

Recent structural studies of 5HT_2A_R and other serotonin receptors identified binding modes of various agonists, inverse-agonists, and antagonists (reviewed in Duan et al.^11^ and Simon et al.^22^) as well as conformational changes during activation and transducer-binding.^18,23–26^ The binding pocket of 5HT_2A_R consists of two adjacent subpockets: the OBP and the extended binding pocket (EBP). Regardless of subpocket localization, ligands are universally anchored in the binding pocket by the central D155^3.32^ residue through interactions with a charged amine moiety.^23,24,27–29^ Additional ligand groups are typically coordinated through hydrophobic interactions and hydrogen bonds by residues in the OBP and EBP. Crystal structures of 5HT_2A_R in complex with serotonin and psilocin showed that the indole core is exclusively located in the EBP, while the OBP and side-extended pocket (SEP) are occupied by a lipid molecule.^28^ On the contrary, computational studies on 5HT_2A_R^23,30^ as well as structural studies on related 5HTRs^18,25,26^ showed an occupation of the OBP by the indole core of serotonin or psilocin. Mutation studies revealed that disruption of coordination in both subpockets diminishes receptor activity, hinting at two binding poses.^28^ Antagonists typically extend beyond the EBP/OBP and additionally reach into the deep binding pocket, where they prevent necessary receptor rearrangements.^27,28^

Activation of 5HT_2A_R is marked by an extensive outward tilt of TM5, and TM6 accompanied by a smaller inward tilt of TM2 and TM3. This TM repositioning in the intracellular part of 5HT_2A_R opens the transducer binding cavity and allows Gqα or β-arrestin to bind. Besides the large helix movements, 5HT_2A_R activation is also marked by structural changes in several functional motifs along the receptor: the rotation of the toggle switch residue W336^6.48^, movement in the PIF motif (P246^5.50^, I163^3.40^, and F332^6.44^), the breaking of the ionic lock between R173^3.50^ and E318^6.30^ as well as the inward shift of the NPXXY motif (N376^7.49^, P377^7.50^, and Y380^7.53^) accompanied by the breaking of the π-stacking interactions between Y380^7.53^ and Y387^8.50.23,24^

In this contribution, we use molecular dynamics (MD) simulations and free energy calculations to delve into the details of the binding of psilocin and serotonin to the two different binding pockets and find that both ligands bind to the OBP with higher affinity. We also investigate the conformational transitions of the receptor and find that the active state is only stable with bound Gqα. We also discover a new conformation where the transitions described in the previous paragraph remain incomplete. We call this the partially-open state of the receptor.

## RESULTS

### ‘Open’ state 5HT_2A_R rapidly relaxes in the absence of Gq*α*

Recent structural studies of 5HT_2A_R produced multiple conformations during different stages of receptor activation. The structures of 5HT_2A_R in an active G protein-bound ‘open’ state as well as in an inactive risperidone-bound ‘closed’ state identified prominent changes in several structural motifs during activation. To capture dynamic structural changes between these endpoints of activation and the influence of agonists on these conformational changes, we performed long all-atom MD simulations using the CHARMM36 force field. We started simulations from both the inactive ‘closed’ and active ‘open’ states and tested the influence of ligands by inserting serotonin and psilocin into the respective subpocket location.

During production, all systems are comparatively stable over time but show a high deviation from the initial structure with root-mean-square distance (RMSD) values of 5-10 Å, independent of the presence of serotonin or psilocin bound the extracellular ligand binding pocket (Figure S1a). Root-mean-square fluctuations (RMSF) analysis shows that this deviation mostly arises from changes in the highly flexible intrinsically disordered intracellular loops (ICLs) and extracellular loops ECLs (Figure S1b) and does not necessarily indicate regulatory conformational changes in the receptor. To capture structural changes in the intracellular transducer binding cavity during transitions from the experimental ‘open’ and ‘closed’ states, we monitored distances between the Cα atoms of R173^3.50^/E318^6.30^ and Q262^5.66^/E318^6.30^ (Figure 2) – assessing the outward tilt of TM5 and TM6. None of the inactive ‘closed’ systems show an opening of the transducer binding cavity via outward movements of TM5 and TM6 as the distance remains at the level of the ‘closed’ experimental reference.^27^ On the other hand, the outward-tilted TM5 and TM6 in the activated ‘open’ state models rapidly collapse to an inward-tilted TM5/TM6 orientation similar to the ‘closed’ state during equilibration or within the first 250 ns of production. Only during one simulation replica did the receptor remain in the ‘open’ state for the full simulation of 1 µs (Figure 2F, blue). To test whether the helix conformation of the ‘open’ state is stable in the presence of a transducer, we performed the same set of MD simulations and intramolecular distance analysis with a full Gqα subunit bound to the ‘open’ intracellular transducer binding cavity and in direct proximity to the ‘closed’ intracellular transducer binding cavity (Figure S2). Except for one replica, the outward orientation of TM5 and TM6 in the ‘open’ state is preserved and in good agreement with the experimental references, while the ‘closed’ systems maintain the inward orientation of TM5 and TM6. The rapid collapse of the ‘open’ state of 5HT_2A_R during the simulations could imply that the large outward movements necessary for adopting the active state seen in structural studies^23,24^ depend on the presence of a bound transducer. Therefore, the dichotomy of an active and inactive state likely represents an incomplete conformational landscape of 5HT_2A_R activation. Before the final binding of a transducer, 5HT_2A_R probably either adopts an intermediate state in a local energy minimum or undergoes a gradual opening of the intracellular binding cavity without adopting a distinct intermediate conformation as seen for other GPCRs.^31^

**Figure 1.**
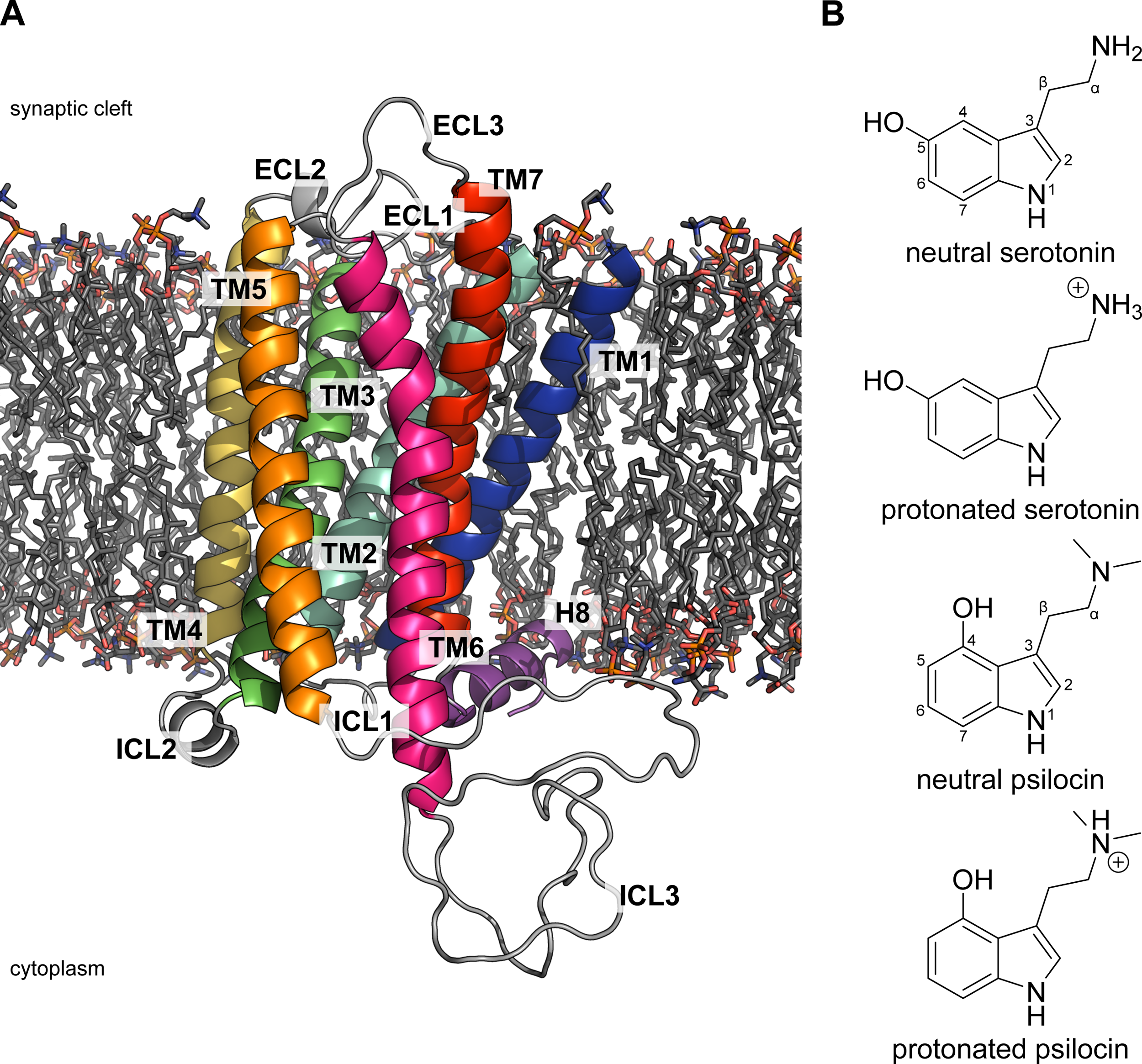
Structures of the simulation model system and ligands. (A) Structure of a 5HT_2A_R (70-399) model inserted into the simplified postsynaptic neural membrane. 5HT_2A_R has a typical GPCR structure with 7 transmembrane helixes connected by intrinsically disordered extracellular and intracellular loops. Water molecules and ions are omitted for clarity. (B) Chemical structure of serotonin and psilocin. The basic _α_-amine group of serotonin (p*K*_a_ 9.97) and psilocin (p*K*_a_ 8.47) can be protonated under physiological conditions.^53^

**Figure 2.**
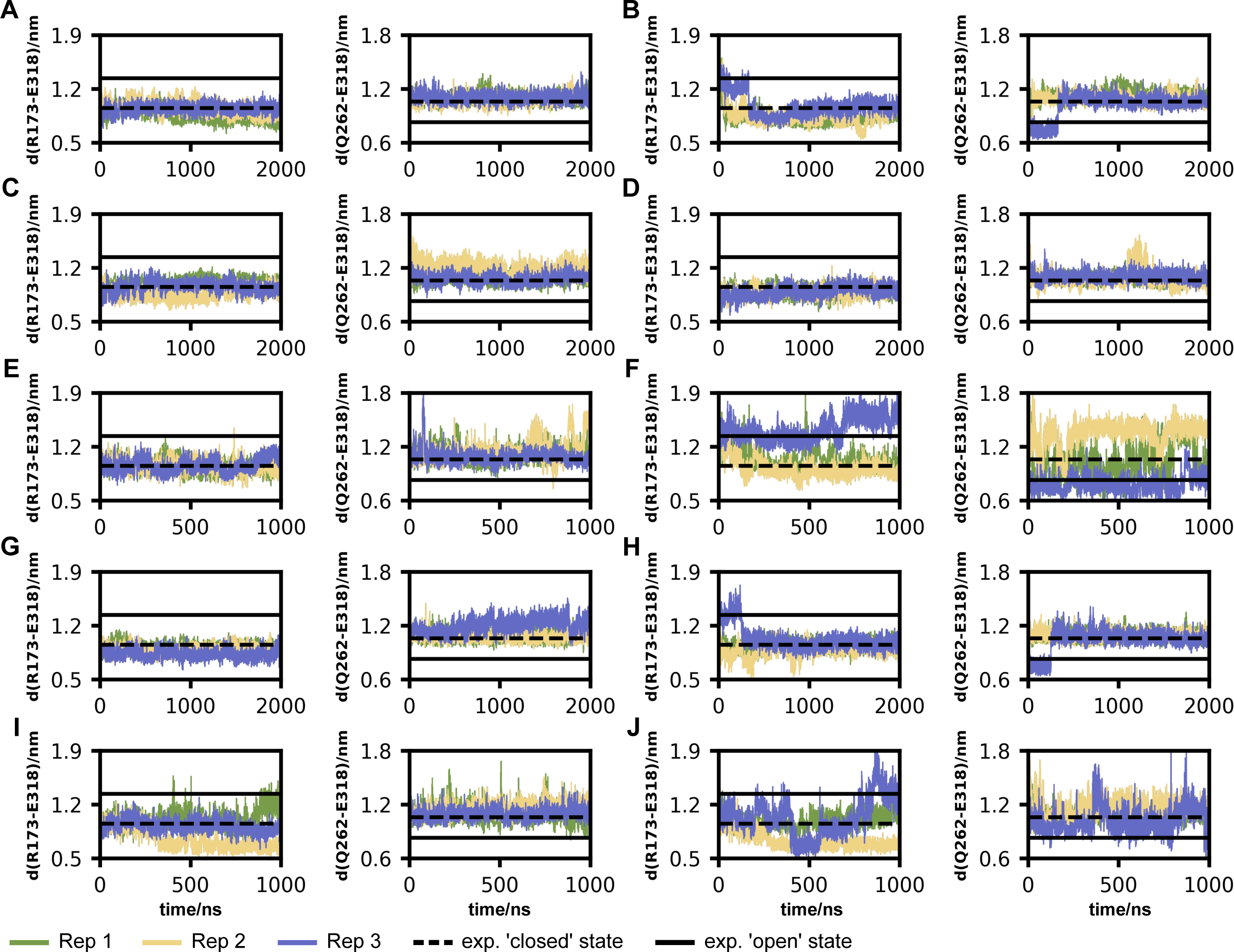
Changes in intramolecular Cα distances characterizing the opening of the intracellular transducer binding cavity through the outward shift of TM5 and TM6 during receptor activation throughout the simulation. (A) ‘Closed’ 5HT_2A_R model without ligand. (B) ‘Open’ 5HT_2A_R model without ligand. (C) ‘Closed’ 5HT_2A_R model with psilocin in OBP. (D) ‘Open’ 5HT_2A_R model with psilocin in OBP. (E) ‘Closed’ 5HT_2A_R model with psilocin in EBP. (F) ‘Open’ 5HT_2A_R model with psilocin in EBP. (G) ‘Closed’ 5HT_2A_R model with serotonin in OBP. (H) ‘Open’ 5HT_2A_R model with serotonin in OBP. (I) ‘Closed’ 5HT_2A_R model with serotonin in EBP. (J) ‘Open’ 5HT_2A_R model with serotonin in EBP. (A–J) The R173-D318 distance also characterizes the breaking of the ionic lock. The horizontal dashed and solid lines are the experimental references for the inactive ‘closed’^27^ and active ‘open’^23,24^ states. Distances in the presence of Gqα are depicted in Figure S2.

### 5HT2AR adopts ‘partially-open’ states during activation

To determine more subtle conformational changes and protein motions in the transmembrane domain of 5HT_2A_R, we performed principal component analysis (PCA) on the backbone atoms of the TM1-7/H8 core. Since the intrinsically disordered connecting loops are much more flexible than the TM1-7/H8 core, and would otherwise dominate the eigenvalue spectrum, ICLs and ECLs were omitted for PCA. For each replica of the systems, up to four conformational ensembles were assigned using *k*-means clustering, and representative conformations were aligned with the experimental ‘closed’ and ‘open’ state structures of 5HT_2A_R (Figure S3).^23,24,27^ Unsurprisingly, the most prominent protein movement occurs in the intracellular short helix H8, which is connected to TM7 only via a short flexible linker and stabilized through π-stacking interactions between Y380^7.53^ and Y387^8.50^. Despite the conformational diversity in this region, none of the ensembles exhibit an inward shift in the NPXXY motif comparable to the Gq-bound experimental structures^23,24^ and are mostly conserved or even strengthened. As seen in the intramolecular distance analysis, the PCA shows that the active ‘open’ systems predominantly adopt a ‘closed’ transducer binding cavity conformation in the absence of Gqα, and the inactive ‘closed’ systems retain their ‘closed’ transducer binding cavity. However, in ∼55 % of the simulations, a small subsection of the conformational ensembles adopts a ‘partially-open’ state with a less extensive outward tilt of TM6 of ∼4 Å instead of the ∼8 Å in the experimental transducer-bound ‘open’ state (Figure 3A).^23,24^ These ‘partially-open’ states could represent an active conformation before Gqα binding that undergoes further opening of the intracellular binding cavity induced by proximity of a transducer molecule. Although the partial outward movement is reminiscent of the activated ‘open’ state of the receptor, the ‘partially-open’ conformations lack other molecular characteristics of activation. Only two of the partially open conformational states undergo a rotation of the toggle switch residue W336^6.48^ and there are no subsequent rearrangements of F332^6.44^ in the PIF motif (Figure 3B). Moreover, the ionic lock between R178^3.55^ and E318^6.30^ is loosened and not fully broken in most conformations. While the salt bridge between the two residues is broken, the side chains remain in comparatively close proximity, so that a reformation of the ionic lock is likely (Figure 3C). Notably, the partial opening of the transducer binding cavity occurs in the absence of ligands as well as in the presence of both serotonin and psilocin. A recent study on G protein activation through the β_2_-adrenergic receptor found a similar ‘partially-open’ intermediate state for the related GPCR by combining time-resolved cryo-electron microscopy and MD simulations.^31^

**Figure 3.**
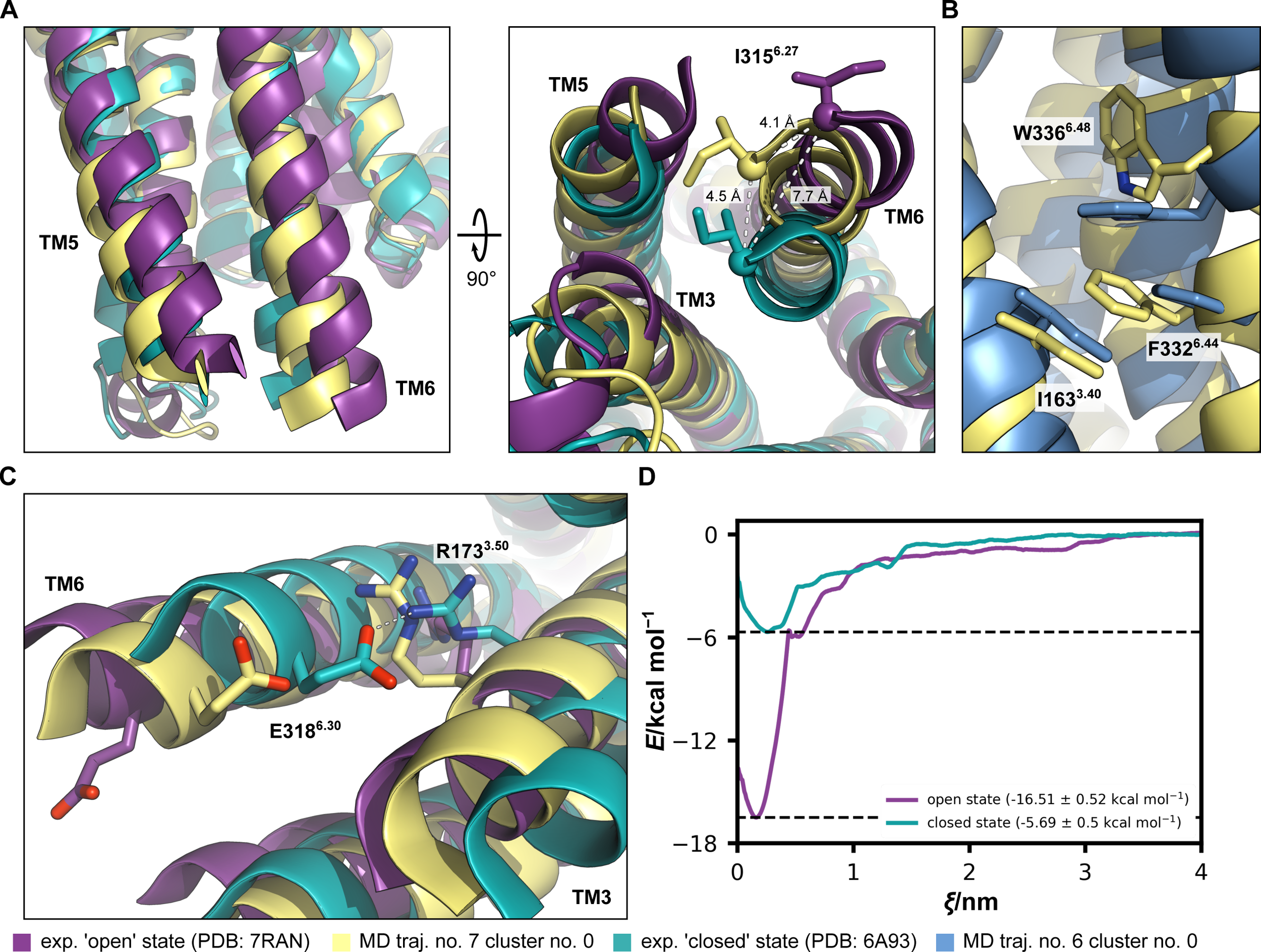
Comparison between the experimental ‘open’/Gq-bound, the ‘partially-open’, and the experimental ‘closed’ states of 5HT_2A_R. (A) Outwards movement of TM6. (B) Rotation of the toggle-switch residue W336^6.48^ and the rearrangement of F332^6.44^ in the PIF motif. (C) Partial breaking of the ionic lock between R173^3.50^ and E318^6.30^. (A–C) An overview of all PCA clusters is depicted in Figure S3. (D) PMF profiles for determining the binding free energy of the C-terminal helix of Gq_α_ to the intracellular transducer binding cavity. Profiles show the detachment of the C-terminal helix of Gq_α_ into bulk water for both the experimental active ‘open’ and inactive ‘closed’ states. For simplicity, the free energy of the system is considered zero when ligands are in bulk water. Data are represented as mean ± SD. Histogram overlaps are depicted in Figure S4.

To investigate the thermodynamic implications of the different conformational states on Gq_α_ binding, we employed potential of mean force calculations to determine the binding affinity of the C-terminal helix of Gqα to the intracellular binding cavity of 5HT_2A_R in the inactive ‘closed’ and the active ‘open’ states (Figure 3D). The binding affinity of the C-terminal Gqα helix is substantially higher to the ‘open’ conformation than to the ‘closed’ conformation. Despite the rapid collapse of the ‘open’ state in the absence of a bound transducer, this difference in binding affinity demonstrates the necessity of a ‘fully-open’ transducer binding cavity for effective Gqα binding. However, the C-terminal Gqα helix still shows a sizeable affinity to the ‘closed’ binding cavity of 5HT_2A_R with a binding free energy of ∼−10.8 kcal mol^−1^. The initial low-affinity binding may serve to align and tether both the receptor and transducer, facilitating the progression toward the subsequent complete opening of the transducer binding cavity, driven by the increasing binding free energy.

### The binding of serotonin and psilocin to the OBP is energetically favored over binding to the EBP

The extracellular ligand binding pocket of 5HTRs consists of the upper EBP as well as the adjacent deeper OBP and is closed through a lid formed by ICL2. Due to the conserved G238^5.42^ residue in the 5HT_2_R family, the OBP is further extended in these receptors via a direct connection to the SEP. This extensive binding pocket allows small ligands like serotonin and psilocin to adopt multiple binding modes by occupying the OBP and EBP, respectively.^11,22^ While the molecular basis of serotonin and psilocin binding has been investigated through structural, computational, and mutational studies,^23,28,30^ the underlying thermodynamic properties and dynamic processes between the two subpockets along with their physiological implications remain unknown.

Thus, we performed potential of mean force calculations to determine the binding affinity of serotonin and psilocin to the two subpockets and compared the affinities between the experimental ‘closed’ and ‘open’ transducer-bound states (Figure 4). For both psilocin and serotonin, the binding free energy differs by [5 kcal mol^−1^ between the OBP and EBP (Table 1), hinting at a more stable binding mode in the deeper OBP as well as a higher relative occupancy of the OBP by psilocin and serotonin. Considering the probability distribution between the two binding modes using a simple Boltzmann factor derived from the differences in the binding free energies, the probability of an occupied OBP is higher by 3 orders of magnitude for the ‘open’ state and 4 orders of magnitude for the ‘closed’ state, respectively. During our conventional MD simulations, the ligands remained stably coordinated in the OBP throughout the full simulations, with ligands escaping the binding pocket in only 2 out of 12 simulations. When placed in the EBP, ligands exhibit more dynamic coordination with movements towards the lid and escape the binding pocket in 5 out of 12 simulations, further underlining the weaker affinity to this subpocket orientation. In the crystal structures of 5HT_2A_R, the occupation of the EBP is accompanied by the occupation of the OBP and SEP through a small monoolein lipid.^28^ This occupation of the OBP/SEP through small lipids gives a molecular explanation for the observation that lipids act as partial agonists^28^ and modulate 5HT_2A_R signaling^32,33^. Our findings suggest that lipids like monoolein, oleamide, or 2-oleoyl glycerol not only modulate 5HT_2A_R activity through competitive binding to the OBP but might be necessary to overcome the energy difference between the two binding modes for effective EBP-mediated signaling.

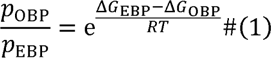

**Figure 4.**
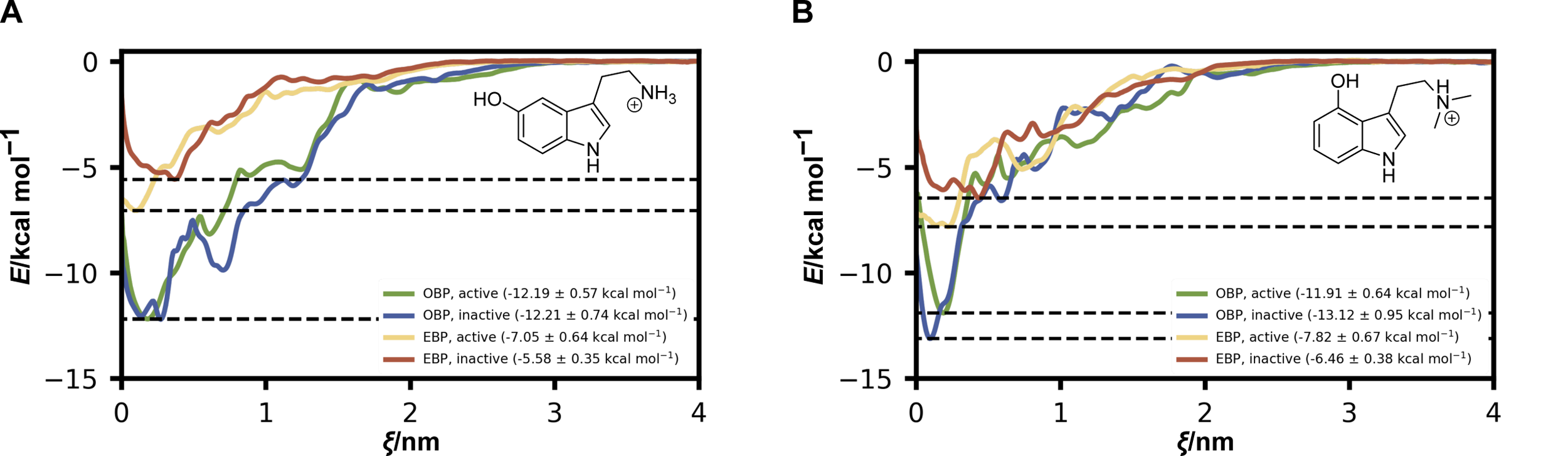
PMF profiles for the determination of the binding free energy of the agonists serotonin and psilocin to the extracellular ligand binding pocket. (A) Profiles show the detachment of serotonin from the OBP and EBP into bulk water for both the experimental active ‘open’ and inactive ‘closed’ states. Data are represented as mean ± SD. Histogram overlaps are depicted in Figure S5. (B) Profiles show the detachment of psilocin from the OBP and EBP into bulk water for both the experimental active ‘open’ and inactive ‘closed’ states. Data are represented as mean ± SD. Histogram overlaps are depicted in Figure S6. (A and B) For simplicity, the free energy of the system is considered zero, when ligands are in bulk water.

**Table 1.** Binding free energies of psilocin and serotonin to the OBP and the EBP.

| System | $\Delta G_{\text{OBP}}/\text{kcal mol}^{-1}$ | $\Delta G_{\text{EBP}}/\text{kcal mol}^{-1}$ | $\Delta\Delta G_{\text{EBP-OBP}}/\text{kcal mol}^{-1}$ |
| --- | --- | --- | --- |
| SERO/open | $-12.19\pm 0.74$ | $-7.05\pm 0.64$ | $5.14\pm 1.21$ |
| SERO/closed | $-12.21\pm 0.74$ | $-5.58\pm 0.35$ | $6.63\pm 1.09$ |
| PSIL/open | $-11.91\pm 0.64$ | $-7.82\pm 0.67$ | $4.09\pm 1.31$ |
| PSIL/closed | $-13.12\pm 0.95$ | $-6.46\pm 0.38$ | $6.66\pm 1.33$ |

Interestingly, there is no substantial difference in binding free energy between the ‘closed’ and ‘open’ states for the OBP. On the other hand, the lower binding free energy and the shift in the reaction coordinate of the energy minimum for the EBP in the ‘closed’ state indicate that both ligands show a higher affinity to the EBP for the receptor in the activated ‘open’ state. The binding of ligands to the EBP during the early stages of receptor activation, when the receptor still adapts a ‘closed’ state, could induce conformational changes towards a more ‘open’ state to accommodate for the lower binding free energy of the ‘closed’ state. Moreover, the lower binding free energy of the EBP magnifies the difference in binding free energy between the two subpockets and therefore shifts the occupation of the two subpockets further towards the OBP for the ‘closed’ state of the receptor. With this magnified difference for the inactive ‘closed’ state, the occupation of the OBP by a lipid molecule might be particularly crucial for EBP-mediated signaling during the early steps of receptor activation.

## DISCUSSION

Using molecular dynamics simulations and free energy calculations, we investigated the binding dynamics of serotonin and psilocin to the 5HT_2A_R and the subsequent conformational changes during receptor activation. Our findings reveal several critical insights into the interaction mechanisms and structural transitions of 5HT_2A_R: Serotonin and psilocin exhibit a higher binding affinity for the OBP over the EBP, which demonstrates stable ligand coordination in the OBP. In the absence of the Gqα subunit, 5HT_2A_R partly adopts a ‘partially-open’ state, characterized by less extensive outward movements of transmembrane helices compared to the fully ‘open’ state observed with Gqα binding. This suggests that the receptor transitions through intermediate states essential for full activation. The active ‘open’ state of 5HT_2A_R demonstrates a significantly higher affinity for the Gqα subunit compared to the ‘closed’ state, emphasizing the importance of a fully open transducer binding cavity for effective Gqα interaction. These findings provide a more detailed mechanistic understanding of the binding dynamics and activation process of 5HT_2A_R, highlighting the significance of receptor-ligand and receptor-transducer interactions. This understanding could guide the design of more selective and effective therapeutics targeting 5HT_2A_R for the treatment of a range of neuropsychiatric disorders, potentially leading to improved therapeutic outcomes with fewer side effects.

## ACKNOWLEDGMENTS

A.A.H.Z. and H.K. are supported by the Novo Nordisk Foundation (#NNF18OC0034936). H.K. is supported by a Lundbeckfonden Ascending Investigator grant number R344-2020-1023. The simulations were performed on the Novo Nordisk Foundation-funded ROBUST Resource for Biomolecular Simulations #NNF18OC0032608.

## AUTHOR CONTRIBUTIONS

N.V. Investigation, data curation, formal analysis, visualization writing – original draft. A.A.H.Z. Conceptualization, methodology, supervision, data curation, formal analysis, writing – review and editing. H.K. Conceptualization, funding acquisition, supervision, project administration, writing – review and editing.

## STARf*Methods

### RESOURCE AVAILABILITY

#### Lead contact

Further information and requests for resources and reagents should be directed to and will be fulfilled by the lead contact, Himanshu Khandelia.

#### Materials availability

This study did not generate new unique reagents.

#### Data and code availability

- All models, data, and scripts were deposited to the Zenodo repository and can be accessed via the DOI: 10.5281/zenodo.12723328
- Any additional information required to reanalyze the data reported in this paper is available from the lead contact upon request.

## METHOD DETAILS

### System construction and equilibration

Experimental protein models of 5HT_2A_R and the Gqα subunit were taken from the Protein Data Bank^34^ as well as the AlphaFold Protein Structure Database^35,36^ and missing side chain atoms were filled in using PyMOL (Schrödinger, LLC). Unresolved ICLs and ECLs were modeled as disordered loops using MODELLER 10.4^37,38^ according to the canonical amino acid sequence of human 5HT_2A_R (UniProtKB accession number: P28223-1). Final receptor models consist of residues 70-399 and contain TM1-TM7, H8, ECL1-ECL3, and ICL1-ICL3 (Figure 1A). Ligands were placed into the OBP according to previous computational studies of 5HT_2A_R^30^ as well as the orientation of ligands in 5HT_1_Rs^25^ and placed into the EBP according to the experimental structures of 5HT_2A_R.^28^ A detailed overview of all systems is given in Table S1.

Assembled protein systems were oriented using the PPM 3.0 Web Server^39^ and constructed using the Membrane Builder tool of CHARMM-GUI.^40,41^ Protein models were inserted into a heterogeneous lipid bilayer membrane consisting of 400 lipids in total with an extracellular leaflet composition of 30 % POPC (16:0/18:1), 12 % PSPC (16:0/18:0), 9 % PAPC (16:0/20:4), 9 % SDPC (18:0/22:6), 10 % SSM, 30 % CHL and an intracellular leaflet composition of 30 % POPC (16:0/18:1), 12 % PSPC (16:0/18:0), 9 % PAPC (16:0/20:4), 9 % SDPC (18:0/22:6), 20 % POPS (16:0/18:1), 20 % CHL, mirroring a simplified lipid composition of a postsynaptic neural membrane.^42–44^ All systems were solvated with TIP3P water with Van der Waals interactions on hydrogen atoms and neutralized with 150 mM KCl using standard GROMACS tools. Initial box sizes and the number of water molecules for the different systems are listed in Table S1.

Systems were energy minimized using the steepest-descent algorithm to remove steric clashes in the initial model. After minimization, systems were equilibrated for 250 ps in an NVT ensemble and 1625 ps in an NPT ensemble using velocity-rescaling temperature coupling^45^ and semi-isotropic stochastic cell rescaling pressure coupling,^46^ while successively reducing position restraints. All simulations were carried out using GROMACS 2023.3^47^ and the CHARMM36 force field (charmm36-jul2022).^48,50^ Topologies and force field parameters for protonated serotonin, protonated psilocin, and guanosine-5’-diphosphate (GDP) were generated using the CHARMM General Force Field (CGenFF) program and converted to GROMACS format using CHARMM36-feb2021 force field and a python script (cgenff_charmm2gmx_py2_nx1.py) from the MacKerell lab.^51,52^ Charge penalty determination and parameter validation were carried out in previous studies and were adopted for simulations in this study.^53,54^ The basic α-amine group of serotonin and psilocin as well as D155^3.32^ in the ligand binding pocket can be protonated or deprotonated under physiological conditions. Previous simulations showed that the interaction between deprotonated anionic D155^3.32^ and protonated cationic serotonin and psilocin is the most stable configuration.^30^ Thus, all simulations were carried out with deprotonated D155^3.32^ and protonated serotonin and psilocin, respectively (Figure 1B).

### Conventional molecular dynamics simulations

For production, systems without Gqα were simulated for a total of 2 µs and systems with Gqα for a total of 1 µs with a time-step of 2 fs. Three replicas of each were simulated with different initial velocity distributions during the first step of equilibration. Temperature and pressure were kept constant at 310 K and 1 bar using the Nosé–Hoover thermostat^55,56^ and the semi-isotropic Parrinello-Rahman barostat.^57^ The Verlet neighbor search algorithm^58^ was used to update the neighbor list, hydrogen bonds were constrained using the Linear Constraint Solver (LINCS) algorithm,^59^ van der Waals interactions were smoothly switched off between 1.0 and 1.2 nm, and electrostatic interactions were treated with the particle mesh Ewald (PME) method.^60,61^

Trajectories were visually inspected using Visual Molecular Dynamics (VMD).^62^ To monitor extensive conformational changes, the RMSD of the protein backbone and the RMSF of every residue were calculated with respect to the aligned initial structure using standard GROMACS tools. Frames every 0.1 ns were analyzed for RMDS and RMSF calculation. Conformational changes associated with 5HT_2A_R activation were assessed using intramolecular distances and PCA. Distances between Cα atoms of R173/E318 and Q262/E318 were determined throughout the simulations using standard GROMACS tools. Reference distances for the ‘inactive’ and ‘active’ states were derived from the respective experimental structures. For PCA, first, mass-weighted covariance matrices of the 5HT_2A_R backbone were calculated and diagonalized. Afterward, 2D projections of trajectories onto the first and second eigenvectors were calculated using standard GROMACS tools. Due to their large inherent flexibility, ICLs (residues 102-108, 179-189, and 263-314) and ECLs (residues 138-145, 217-231, and 347-354) were omitted for analysis. Centroids of clusters correspond to representative conformational states and were determined using a *k*-means clustering algorithm. For comparison, representative conformational states were aligned with the experimental ‘inactive: (PDB: 6A93)^27^ and ‘active’ (PDB: 7RAN)^24^ state structures of 5HT_2A_R.

### Potential of mean force calculations

Binding free energies of psilocin and serotonin to the OBP and EBP as well as the C-terminal helix of Gqα to the intracellular binding cavity in different states were determined through PMF calculations using umbrella sampling simulations.^63^ The C-terminal helix of Gqα is the only Gqα entity that contacts the receptor. Therefore, the binding free energy between the helix and the receptor is an excellent proxy for the overall receptor-Gqα binding free energy. Simulations of the helix instead of the entire Gqα save significant computational resources. Furthermore, protein-protein binding affinity measurements between large proteins along a distance reaction coordinate can suffer from the influence of other degrees of freedom such as protein rotation which can confound the PMF.^64^ Systems were prepared and equilibrated as described above. Since the TMs of the ‘active’ state model rapidly repositioned during conventional MD simulations, the positions of backbone atoms were restrained with a force constant of 100 kJ mol**^−^**^1^ nm**^−^**^2^. The hydrogen bond network of the C-terminal Gqα helix dissipated partly during simulations of most umbrella windows, so distance restraints between Cα atoms were used to maintain the proper secondary structure of the helix. Initial configurations along the reaction coordinate ξ, defined as the *z*-axis (membrane normal), were created through steered molecular dynamics (SMD) simulations by pulling the ligands from the respective binding site for 500 ps with a force constant of 1000 kJ mol**^−^**^1^ nm**^−^**^2^ and a rate of 0.01 nm ps**^−^**^1^. The resulting 40 windows per simulation had a uniform spacing of ∼0.1 nm over a total of ∼4 nm. Regions with a lack of sampling were complemented with additional asymmetric windows according to the umbrella histograms. Each window was equilibrated for 1 ns and conformations were subsequently sampled for 100 ns/50 ns (ligand binding/Gqα binding), for a total simulation time of ∼4 µs/∼2 µs per system. The initial center of mass (COM) distance between the ligand and the binding site in every window was restrained with a force constant of 1000 kJ mol**^−^**^1^ nm**^−^**^2^. Other simulation parameters were like the parameters of conventional MD simulations described above. PMF profiles were extracted with the Weighted Histogram Analysis Method (WHAM)^65^ and statistical errors were estimated through Bayesian bootstrap analysis^66^ using the GROMACS WHAM module.^67^

## SUPPLEMENTAL INFORMATION

Document S1. Figures S1 – S6 and Table S1

