## Supplemental figures and tables for "Molecular insights into the modulation of the 5HT_2A_ receptor by serotonin, psilocin, and the G protein subunit Gqα"

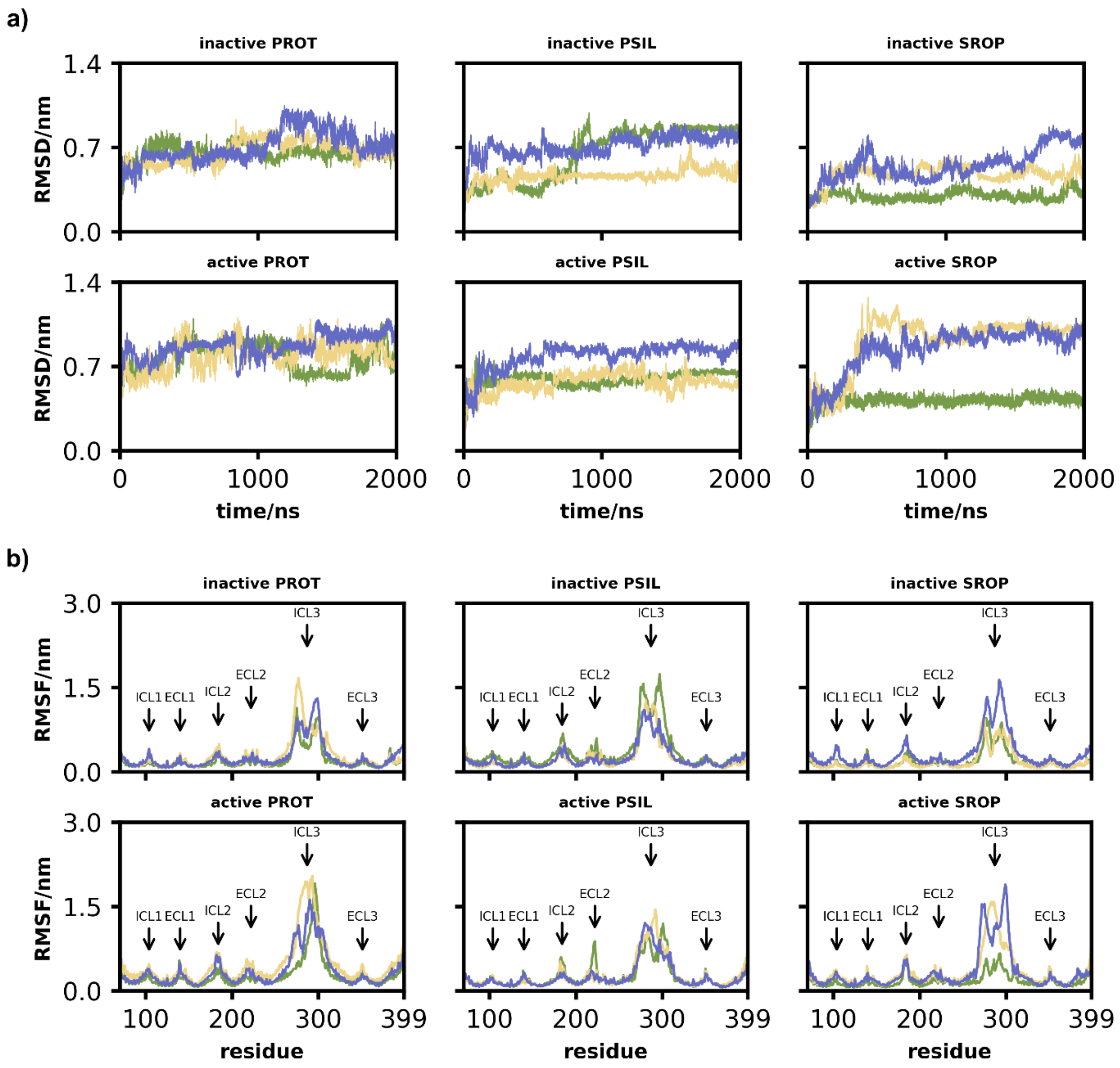


**Figure S1. Conformational changes during simulation of $5HT_{2A}R$.**

(A) RMSD of the $5HT_{2A}R$ backbone from the initial structure throughout the MD simulation. Time points every 0.1 ns were analyzed.

(B) RMSF of $5HT_{2A}R$ residues from the initial structure throughout the MD simulation. The particular flexible loops between the TMs are marked.

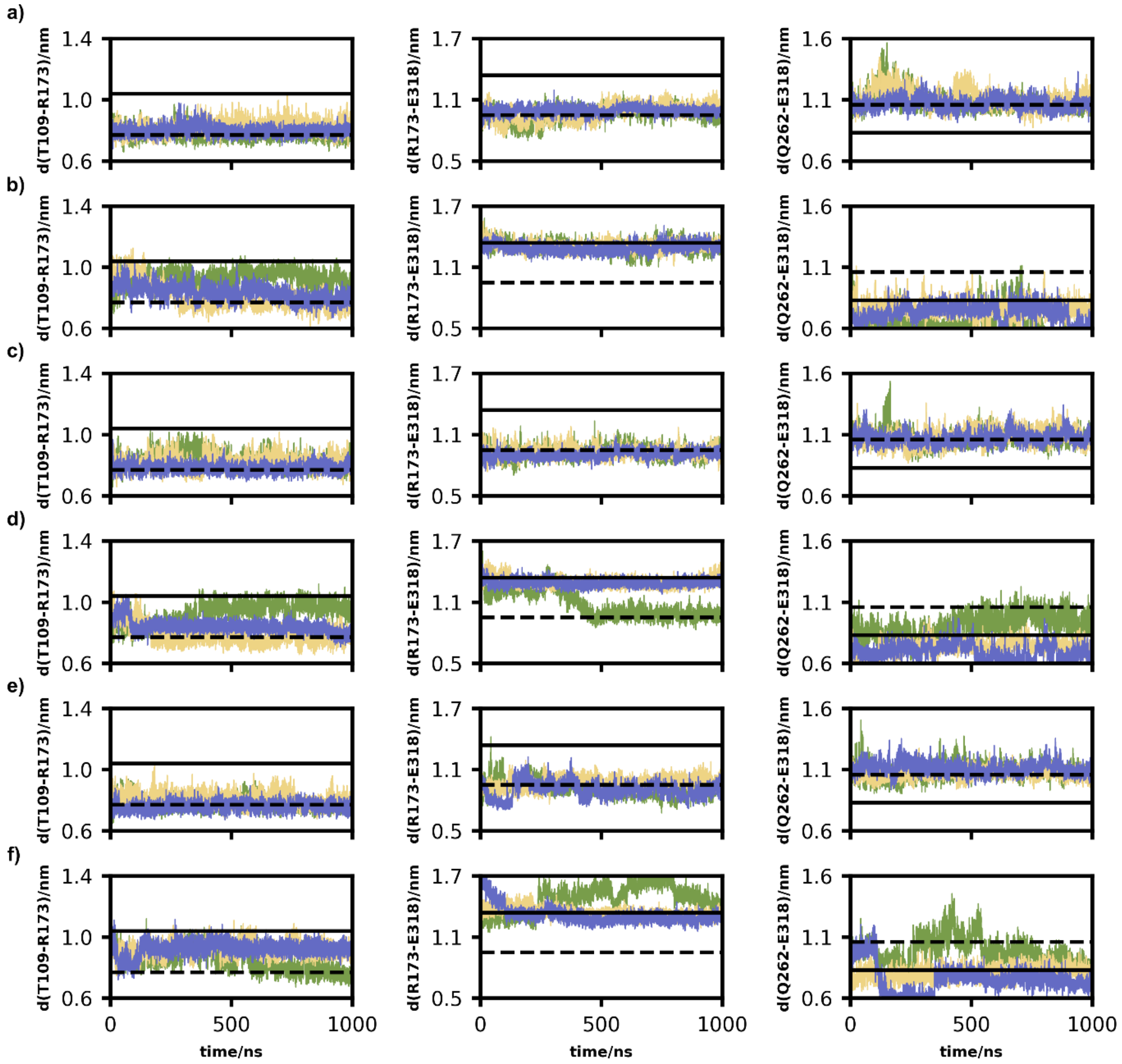


**Figure S2. Changes in intramolecular Cα distances characterizing changes during receptor activation throughout the full $5HT_{2A}R$ simulation in the presence of the transducer subunit Gqα, related to Figure 2.**

(A) 'Inactive' $5HT_{2A}R$ model without ligand.

(B) 'Active' $5HT_{2A}R$ model without ligand.

(C) 'Inactive' $5HT_{2A}R$ model with PSIL in the OBP.

(D) 'Active' $5HT_{2A}R$ model with PSIL in the OBP.

(E) 'Inactive' $5HT_{2A}R$ model with SERO in the OBP.

(F) 'Active' $5HT_{2A}R$ model with SERO in the OBP.

(A–F) The outward shift of TM5 and TM6 is assessed through the R173-E318 distance and Q262-E318 distance, respectively. Additionally, the former also characterizes the breaking of the ionic lock. The inward shift of TM2 and TM3 is measured via the T109-R173. Dashed: Intramolecular distance in the experimental 'inactive' structure. Solid: Intramolecular distance in the experimental 'active' structure.

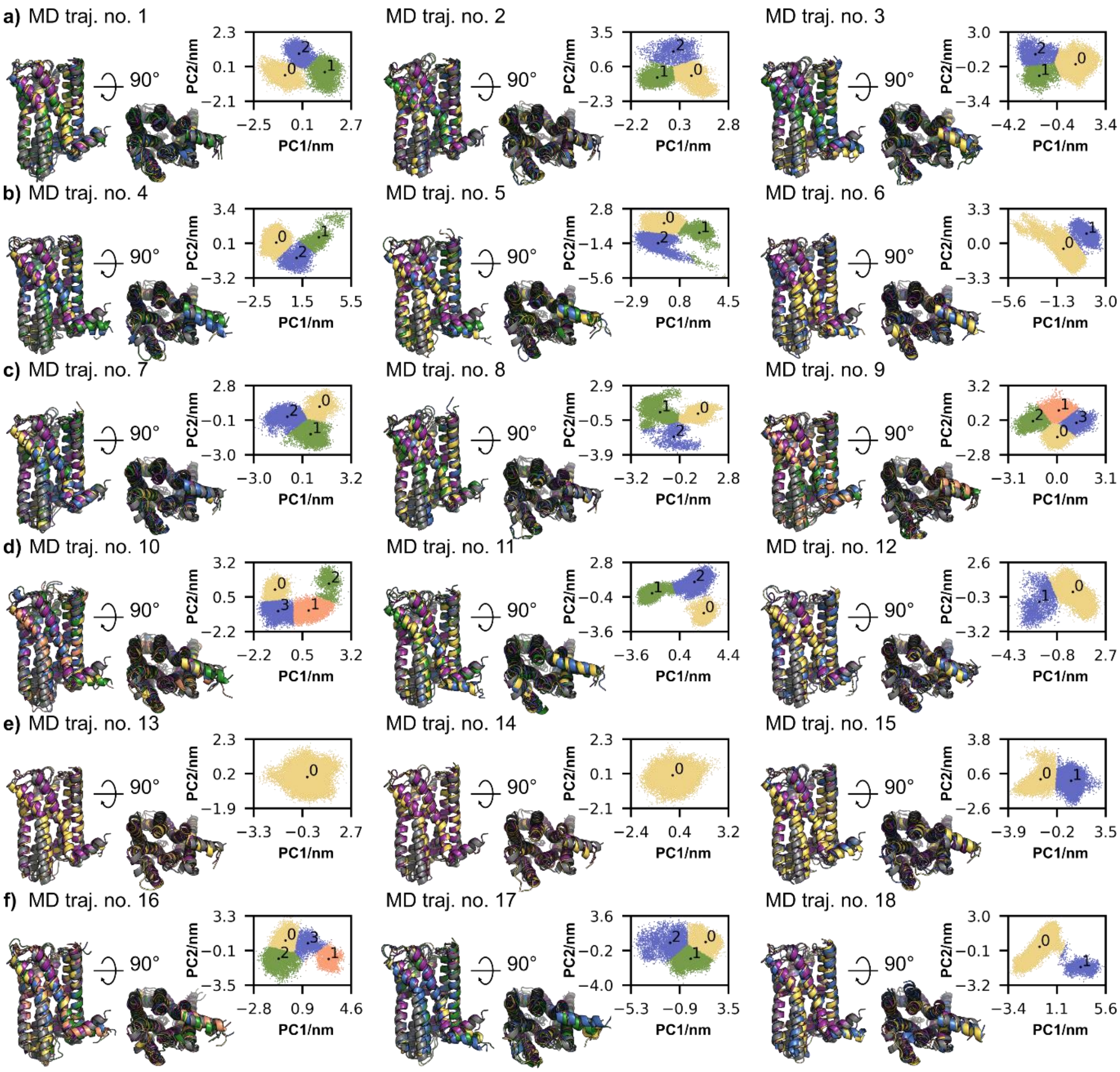


**Figure S3. Principal component analysis of the conformational changes in the 5HT$_{2A}$R backbone during simulation, related to Figure 3A–C.**

(A) ‘Inactive’ 5HT$_{2A}$R model without ligand.

(B) ‘Active’ 5HT$_{2A}$R model without ligand.

(C) ‘Inactive’ 5HT$_{2A}$R model with PSIL in the OBP.

(D) ‘Active’ 5HT$_{2A}$R model with PSIL in the OBP.

(E) ‘Inactive’ 5HT$_{2A}$R model with SERO in the OBP.

(F) ‘Active’ 5HT$_{2A}$R model with SERO in the OBP.

(A–F) 2D projection of the three independent trajectories on each system’s first and second eigenvectors. Due to the high conformational flexibility, ICLs and ECLs were omitted from the analysis. Centroid structures were determined using k-means clustering and aligned to the experimental ‘inactive’ (purple, PDB: 6a93) and ‘active’ (gray, PDB: 7ran) structures.

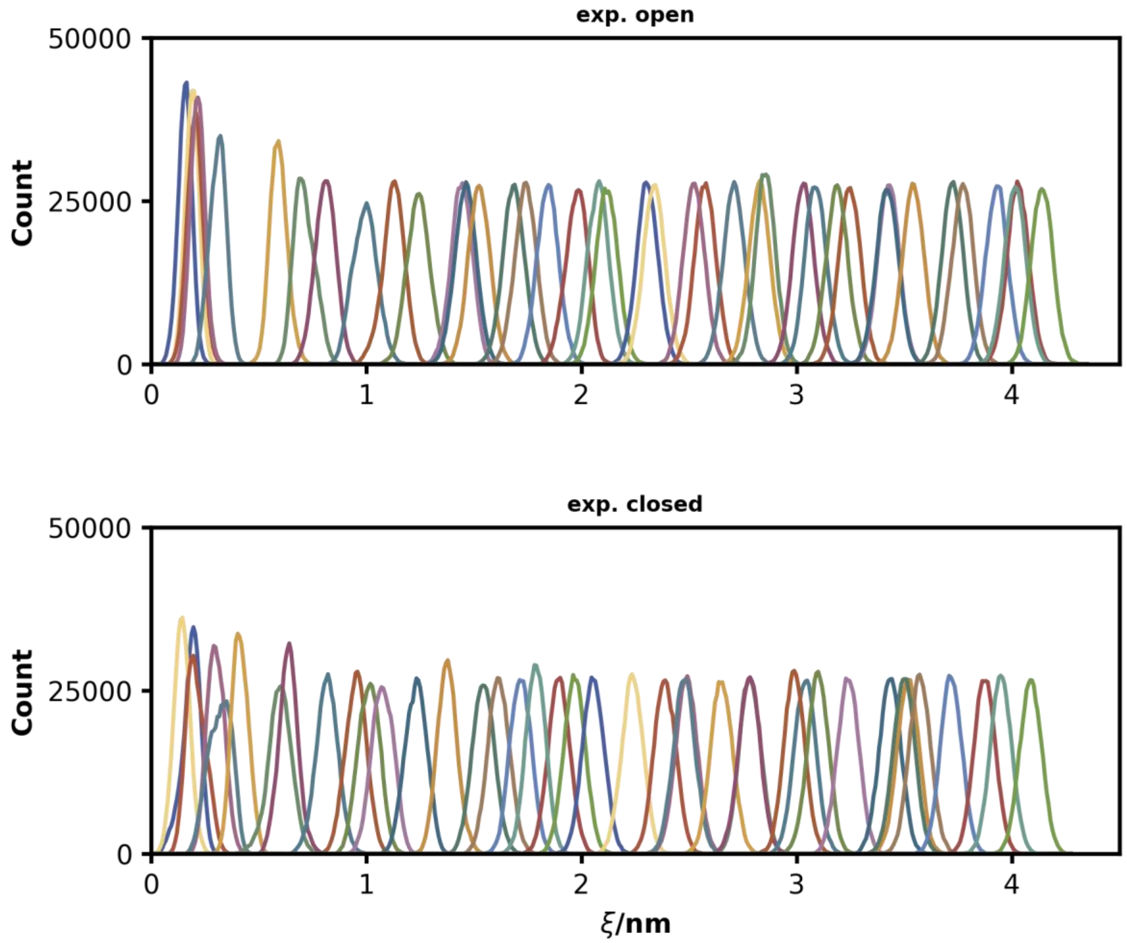


**Figure S4. Umbrella histograms for the PMF calculation of Gqα binding to the intracellular transducer binding cavity of $5HT_{2A}R$, related to Figure 3D.**

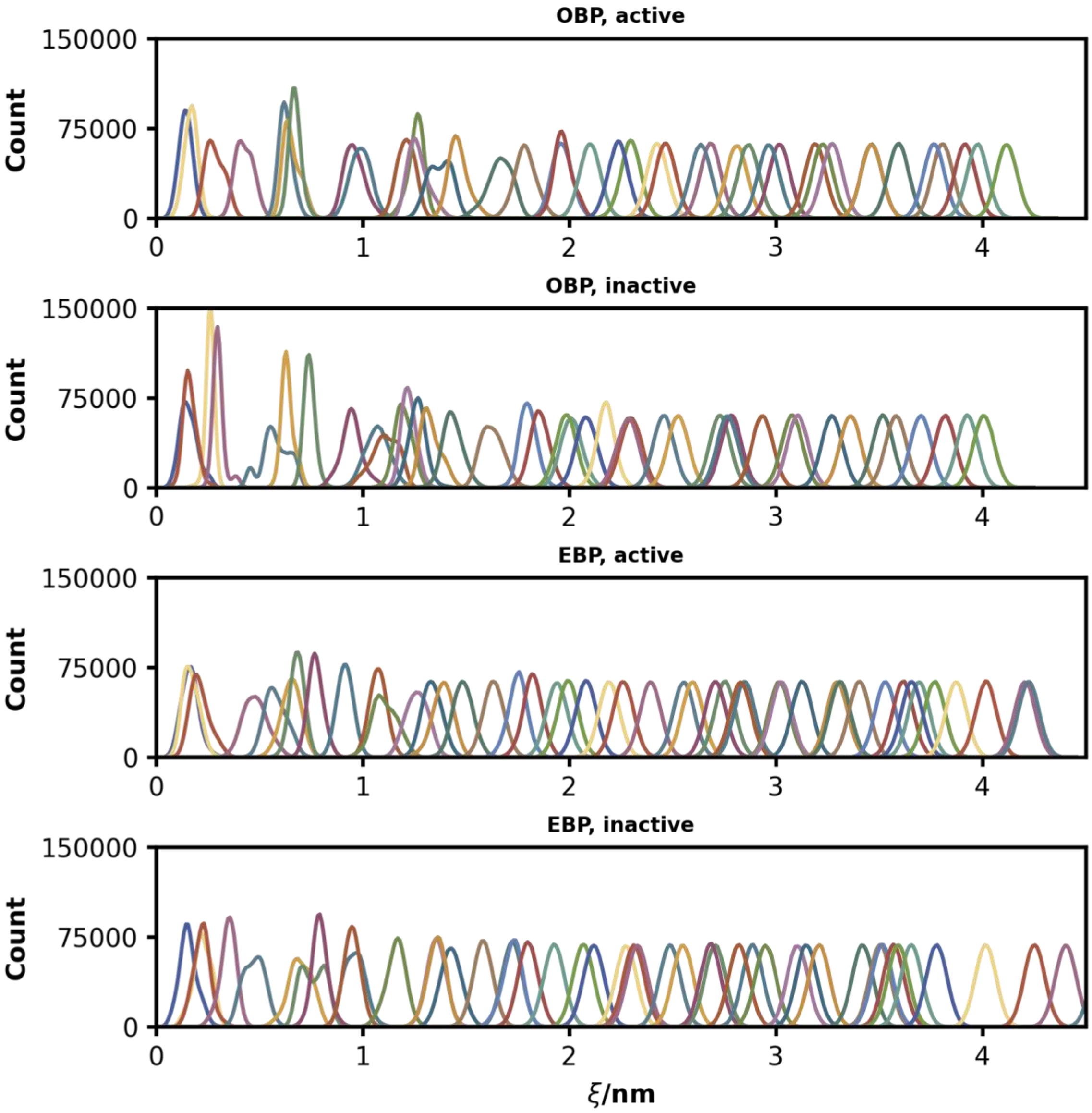


**Figure S5. Umbrella histograms for the PMF calculation of serotonin binding to the extracellular binding pocket of $5HT_{2A}R$, related to Figure 4A.**

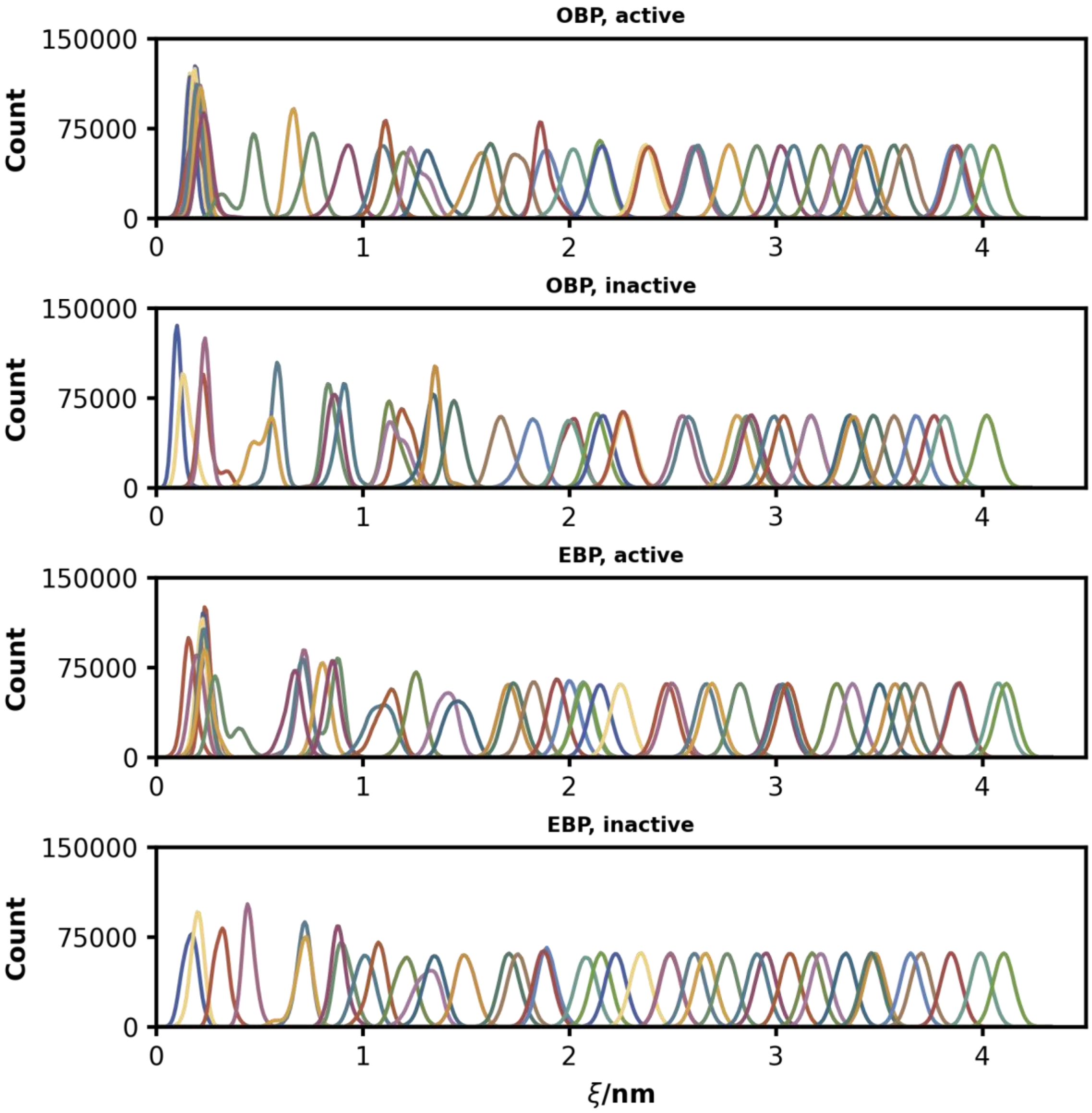


**Figure S6. Umbrella histograms for the PMF calculation of psilocin binding to the extracellular binding pocket of $5HT_{2A}R$, related to Figure 4B.**

**Table S1. Overview of system properties for MD simulations.**

| $5HT_{2A}R$ model | ligand | transducer | box size/Å$^3$ | no. water | no. lipids | traj. no. |
|---|---|---|---|---|---|---|
| *Conventional MD simulations* | | | | | | |
| 7RAN* | — | — | 115×115×147 | ~43000 | 400 | 4, 5, 6 |
| 7RAN* | $SERO^+$ (OBP) | — | 105×105×161 | ~43000 | 400 | 10, 11, 12 |
| 7RAN* | $PSIL^+$ (OBP) | — | 105×105×161 | ~43000 | 400 | 16, 17, 18 |
| 6WHA* | $SERO^+$ (EBP) | — | 115×115×157 | ~53000 | 400 | 22, 23, 24 |
| 6WHA* | $PSIL^+$ (EBP) | — | 115×115×157 | ~53000 | 400 | 28, 29, 30 |
| 6A93† | — | — | 115×115×163 | ~50000 | 400 | 1, 2, 3 |
| 6A93† | $SERO^+$ (OBP) | — | 105×105×183 | ~50000 | 400 | 7, 8, 9 |
| 6A93† | $PSIL^+$ (OBP) | — | 105×105×183 | ~50000 | 400 | 13, 14, 15 |
| 6A93† | $SERO^+$ (EBP) | — | 115×115×157 | ~53000 | 400 | 19, 20, 21 |
| 6A93† | $PSIL^+$ (EBP) | — | 115×115×157 | ~53000 | 400 | 25, 26, 27 |
| 6WHA* | — | Gqα‡ | 115×115×210 | ~74000 | 400 | 34, 35, 36 |
| 6WHA* | $SERO^+$ (OBP) | Gqα‡ | 115×115×210 | ~74000 | 400 | 40, 41, 42 |
| 6WHA* | $PSIL^+$ (OBP) | Gqα‡ | 115×115×210 | ~74000 | 400 | 46, 47, 48 |
| 6A93† | — | Gqα‡ | 115×115×210 | ~74000 | 400 | 31, 32, 33 |
| 6A93† | $SERO^+$ (OBP) | Gqα‡ | 115×115×210 | ~74000 | 400 | 37, 38, 39 |
| 6A93† | $PSIL^+$ (OBP) | Gqα‡ | 115×115×210 | ~74000 | 400 | 43, 44, 45 |
| *PMF calculations* | | | | | | |
| 6WHA** | — | Gqα‡‡ | 115×115×152 | ~50000 | 400 | — |
| 6A93†† | — | Gqα‡‡ | 115×115×152 | ~50000 | 400 | — |
| 6WHA** | — | Gqα‡‡‡ | 115×115×152 | ~50000 | 400 | — |
| 6WHA** | $SERO^+$ (OBP) | — | 115×115×157 | ~53000 | 400 | — |
| 6WHA** | $PSIL^+$ (OBP) | — | 115×115×157 | ~53000 | 400 | — |
| 6WHA** | $SERO^+$ (EBP) | — | 115×115×157 | ~53000 | 400 | — |
| 6WHA** | $PSIL^+$ (EBP) | — | 115×115×157 | ~53000 | 400 | — |
| 6A93†† | $SERO^+$ (OBP) | — | 115×115×157 | ~53000 | 400 | — |
| 6A93†† | $PSIL^+$ (OBP) | — | 115×115×157 | ~53000 | 400 | — |
| 6A93†† | $SERO^+$ (EBP) | — | 115×115×157 | ~53000 | 400 | — |
| 6A93†† | $PSIL^+$ (EBP) | — | 115×115×157 | ~53000 | 400 | — |

* ‘open’/Gqα-bound state. ** ‘open’/Gqα-bound state with position restraints on $5HT_{2A}R$ backbone atoms. † ‘closed’ state. †† ‘closed’ state with position restraints on $5HT_{2A}R$ backbone atoms. ‡ AlphaFold prediction (res 335-359 were replaced with PDB: 6WHA and GDP was placed according to PDB: 7W40). ‡‡ res 335-359 from PDB: 6WHA with distance restraints on Cα atoms. ‡‡‡ AlphaFold prediction without res 335-359 (GDP was placed according to PDB: 7W40) with distance restraints on Cα atoms.
